# Ontogeny of the VIP+ interneuron sensory-motor circuit prior to active whisking

**DOI:** 10.1101/2020.07.01.182238

**Authors:** Cristiana Vagnoni, Liad J. Baruchin, Filippo Ghezzi, Sara Ratti, Zoltán Molnár, Simon J. B. Butt

## Abstract

Development of the cortical circuits for sensory-motor processing require the coordinated integration of both columnar and long-range synaptic connections. To understand how this occurs at the level of individual neurons we have explored the timeline over which vasoactive intestinal peptide (VIP)-expressing interneurons integrate into mouse somatosensory cortex. We find a distinction in emergent long-range anterior-motor and columnar glutamatergic inputs onto layer (L)2 and L3 VIP+ interneurons respectively. In parallel, VIP+ interneurons form efferent connections onto both pyramidal cells and interneurons in the immediate column in an inside-out manner. Cell-autonomous deletion of the fate-determinant transcription factor, *Prox1*, spares long-range anterior-motor inputs onto VIP+ interneurons, but leads to deficits in local connectivity. This imbalance in the somatosensory circuit results in altered spontaneous and sensory-evoked cortical activity *in vivo*. This identifies a critical role for VIP+ interneurons, and more broadly interneuron heterogeneity, in formative circuits of neocortex.

## INTRODUCTION

The mammalian neocortex is an exquisitely assembled neural circuit for higher cognitive functions such as the detection of changes in the environment and initiation of voluntary movement. Over recent years, advances in technology have afforded us a greater understanding of how information is encoded across the different cortical areas giving rise to brain-wide information processing and behavioural output at juvenile and adult stages (Barson et al., 2020; Xiao et al., 2017). However to date developmental studies have largely focused on emergent connectivity within a given sensory area (e.g. Erzurumlu and Gaspar, 2012; Hensch, 2005) and, as a consequence, relatively little is known about how long range synaptic connections integrate with local circuits within the first few postnatal weeks (Arruda-Carvalho et al., 2017; De León Reyes et al., 2019). The need for such an understanding is evident in somatosensory barrel field (S1BF), a primary sensory neocortical area that requires processing of both motor – active exploration of the target by the vibrissae – and tactile sensory information arriving in S1BF (Petersen, 2019).

To specifically explore the emergence of long-range and local connections in S1BF development, we focused on a subtype of GABAergic interneuron defined by the expression of vasoactive intestinal peptide (VIP) (Demeulemeester et al., 1988; Hendry et al., 1984; Kawaguchi and Kubota, 1996). VIP+ interneurons (INs) represent one of the major subtypes of 5-HT_3A_R-expressing GABAergic interneuron (Lee et al., 2010; Rudy et al., 2011) that originate from the ventricular zone of the caudal ganglionic eminence (CGE) in the embryonic telencephalon. In line with other CGE-derived INs, VIP+ INs preferentially populate supragranular layers of neocortex within the first postnatal week (Miyoshi and Fishell, 2011) under the control of serotonin (5-HT) (Frazer et al., 2015), the transcription factor *Prox1* (Miyoshi et al., 2015), and local cues from pyramidal cells (Wester et al., 2019). Three lines of evidence suggest that VIP+ INs represent a key neuronal population that orchestrates sensory-motor processing and a valid target for our investigation: first, VIP+ INs have been shown to play a key role in integrating local and long range inputs in the mature brain (Lee et al., 2013; Wall et al., 2016; Zhang et al., 2014), despite only comprising a small fraction of the total neuronal population in neocortex (Rudy et al., 2011; Xu et al., 2010). Second, VIP+ INs are thought to control information processing in cortical circuits primarily through disinhibition of another class of GABAergic interneuron defined by expression of the neuropeptide somatostatin (SST) (Lee et al., 2013; Pfeffer et al., 2013; Pi et al., 2013). SST+ INs play a prominent role in development, directing synapse formation (Oh et al., 2016), circuit maturation (Tuncdemir et al., 2016), and sensory integration (Marques-Smith et al., 2016). Third, dysfunction of VIP+ INs is found in models of neurodevelopmental psychiatric disorders pointing to an important role for this type of interneuron in early development (Batista-Brito et al., 2017; Goff and Goldberg, 2019; Mossner et al., 2020).

Our aim was to better understand the timeline over which VIP+ INs integrate into both local and long range neocortical circuits in the immediate postnatal time window, expanding on existing knowledge that has identified that CGE-derived INs can receive input from the thalamus and basal forebrain from as early as the first postnatal week (Che et al., 2018; De Marco García et al., 2015). Furthermore we wanted to understand if VIP+ INs influence transient IN circuits (Marques-Smith et al., 2016) and thereby contribute to circuit maturation, before finally probing the requirement for normal VIP+ INs synaptic integration in emergent activity in juvenile neocortex through cell autonomous deletion of *Prox1*.

We focused our investigation on emergent VIP+ IN connectivity and function in mouse whisker barrel cortex (S1BF) as we have a good understanding of key milestones in the development of this sensory modality (Erzurumlu and Gaspar, 2012). Moreover, the distribution of VIP+ INs across the depth of the cortex (Prönneke et al., 2015) and contribution to both columnar and cross-modal integration is well described in adult (Lee et al., 2013; Pfeffer et al., 2013; Pi et al., 2013; Wall et al., 2016; Zhang et al., 2014). Our data reveal that VIP+ INs integrate into cortical circuits as early as the layer 4 critical period for plasticity (CPP), towards the end of the first postnatal week. Moreover we observe a progressive maturation of both local and long-range afferent input prior to the onset of active perception towards the end of the second postnatal week. These observations highlight the importance of VIP+ IN pathway for the establishment of formative networks of the mammalian cerebral cortex for sensory-motor integration.

## RESULTS

### Emergence of vasoactive intestinal peptide (VIP+)-expressing interneurons in early postnatal somatosensory cortex

To label VIP+ INs in neonatal whisker barrel somatosensory (S1BF) cortex, we generated *VIPCre;Ai9* (tdTomato+) offspring and confirmed co-expression of VIP and calretinin in tdTomato+ cells at postnatal day (P)21 (**Supplemental Figure 1a**). Next, we assessed the number and distribution of tdTomato+ cells across the depth of the cortex through postnatal development (P1-P21) (**Figure 1a-c and Supplemental Figure 1 c-d**). This revealed that the density of tdTomato+ cells in S1BF increased over the first postnatal week (**Figure 1b**), with the majority of cells located in superficial cortical layers throughout early development (**Figure 1c**). The low number of tdTomato+ cells at P3 could result from delayed migration of this CGE-derived interneuron subtype (Miyoshi and Fishell, 2011; Miyoshi et al., 2010; Taniguchi et al., 2011) or late conditional expression of the reporter allele. Nevertheless, these data confirmed that the conditional genetic strategy is effective in labelling VIP+ INs from around the end of the first postnatal week onward (Prönneke et al., 2015; Taniguchi et al., 2011). As such, subsequent experiments were targeted to three experimental time windows relevant to emergent perception in S1BF: (1) the critical period of plasticity for layer (L)4 (CPP; defined as P5-P8); (2) the interval following CPP and preceding active whisking (pre-AW; P9-P11); and (3) the period following onset of active whisking (AW; P12-P16).

**Figure 1.**
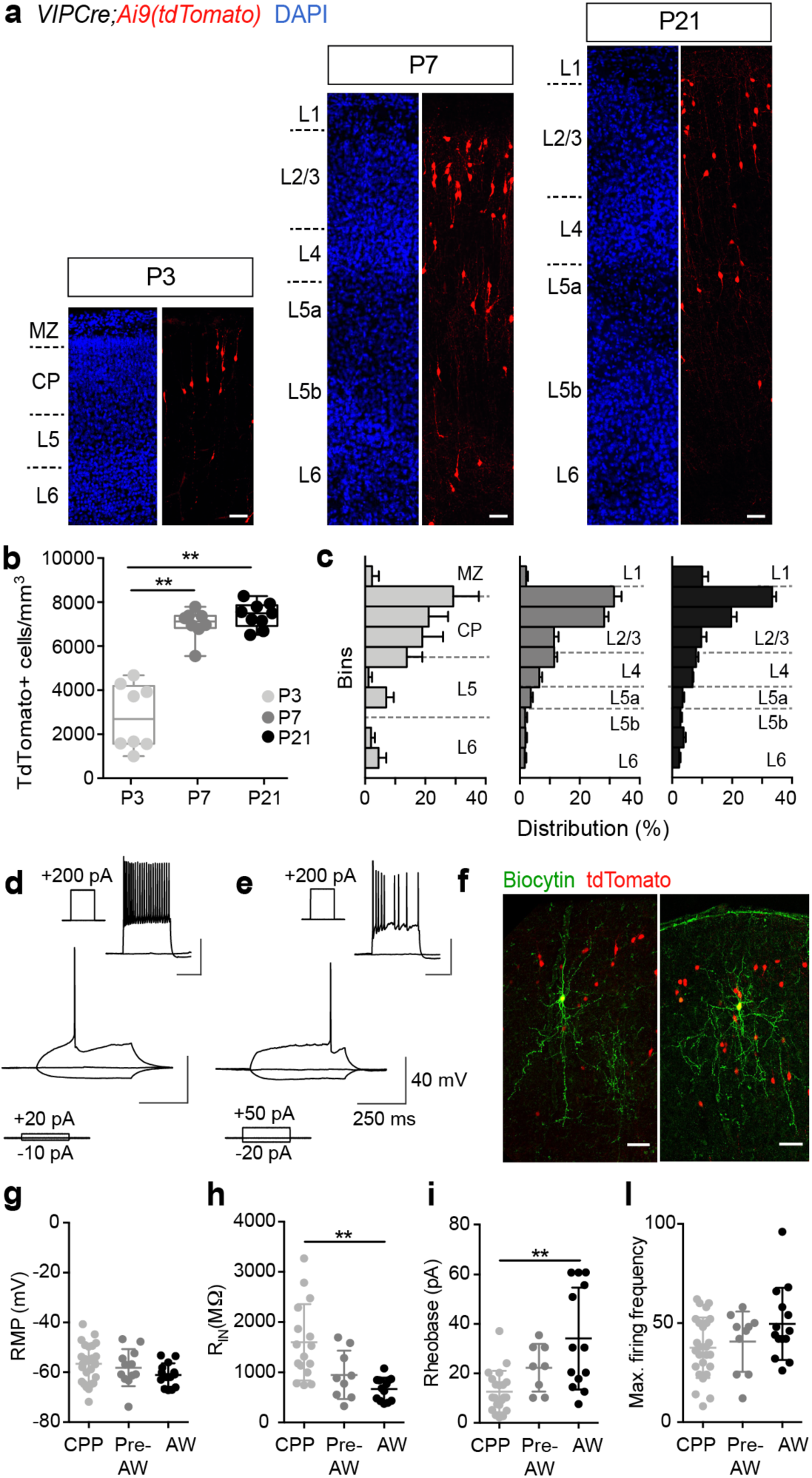
Characterization of S1BF VIP+ INs throughout early postnatal development. (a) Images of S1BF from the *VIP-Cre;Ai9* line at P3, P7 and P21. Blue: DAPI, red: tdTomato. Scale bar: 50 µm. MZ, marginal zone; CP, cortical plate; L, layer. (b) Density of tdTomato+ cells across time points. P3 and P7: 8 counts from n=3 animals; P21: 9 counts from n=3 animals. One way ANOVA: F (2, 22) = 55.44, P< 0.0001; *Post hoc* Tukey’s multiple comparisons test P3 vs. P7: P ≤ 0.0001**, P3 vs. P21: P ≤ 0.0001**. (c) Normalized distribution of tdTomato+ cells (mean ± SEM) across the cortical depth from the pial surface to the white matter tract at each time point (see legend in b). (d,e) Intrinsic electrophysiological profiles of two distinct VIP+ INs subtypes at AW: (d) continuous adapting and (e) irregular spiking interneurons. Superimposed traces show threshold spike, resting membrane potential and response to hyperpolarizing square pulse current injections as indicated in the bottom left corner. Inset: corresponding near maximal firing frequency responses for each subtype; scale bars consistent with the main image. (f) Morphologies of recovered VIP+ INs recorded at AW: bipolar (left) and multipolar (right) subtypes. Scale bar 50 µm. (g-l) Passive and active intrinsic electrophysiological properties recorded in VIP+ INs through development. Horizontal line indicates the mean and error bars the standard deviation. RMP: one-way ANOVA=ns; Max firing frequency: Kruskal-Wallis (K-W) ANOVA=ns; R_IN_ and Rheobase: K-W ANOVA followed by Dunn’s multiple comparisons test. **: P≤ 0.01

To further characterise the tdTomato+ population, we performed whole-cell patch clamp recordings of fluorescent L2/3 cells (n=52) in acute *in vitro* slices of S1BF to assess intrinsic electrophysiological properties as well as cell morphology. At the latest ages examined (AW), we observed heterogeneity in the intrinsic electrophysiological profiles of tdTomato+ cells consistent with previously reported continuously adapting and irregular spiking VIP+ INs (**Figure 1d,e**) (Prönneke et al., 2015, 2018). Recovered morphologies resembled bipolar and multipolar VIP+ interneurons located in L2/3 (**Figure 1f**) (Prönneke et al., 2015, 2018). While passive and active intrinsic electrophysiological properties (**Figure 1g-l**) point to a progressive maturation of VIP+ INs over the time period studied (**Supplementary table 1**), there was some indication of increasing heterogeneity at later ages, for example increased variance in rheobase (**Figure 1i**). These data reveal that our genetic strategy can reliably identify VIP+ INs through the first postnatal weeks and hence can be used to study their emergent connectivity.

### Glutamatergic afferent input reveals emergence of two distinct supragranular VIP+ IN populations through postnatal development

We employed whole-cell patch-clamp electrophysiology in combination with laser scanning photostimulation (LSPS) of caged glutamate to determine the timeline for synaptic integration of L2/3 VIP+ INs into the local cortical circuit. Excitatory postsynaptic currents (EPSCs) were distinguished from direct glutamate responses based on kinetics and onset relative to the UV (355nm) laser pulse (**Figure 2a**). This analysis revealed that VIP+ INs received local L2/3 glutamatergic input at the earliest ages recorded (CPP), but as development continued, a subset of cells acquired translaminar input, primarily from L4 (**Figure 2b,c**). To characterize the observed heterogeneity in glutamatergic input through development (**Figure 2c**), we performed principal component analysis (PCA) followed by k-means cluster analysis on normalized layer inputs for the VIP+ INs across developmental stages (**Supplemental Figure 2a**). K-means analysis (**Supplemental Figure 2b,c**) identified that glutamatergic afferent input on VIP+ INs was best captured by two clusters: the first dominated by local L2/3 input (termed Local) and the second exhibiting a significant L4 (Translaminar) component (L2/3 input: 79±10% for Local vs. 49±11% for Translaminar cluster; L4 input: 8±5% for Local vs. 36±15% in Translaminar)(**Figure 2d,e**). Local VIP+ INs occupied more superficial, L2 locations compared to Translaminar cells across the time period studied (**Figure 2f**). The incidence of translaminar input increased in the Pre-AW and AW time windows (**Figure 2g**; **Supplemental Figure 2a,d**). These data show that VIP+ INs are integrated in the local L2/3 glutamatergic network from as early as the CPP. In addition our results suggest diversity in VIP+ INs based on layer location (L2 *versus* L3) as a result of emergent translaminar feed-forward excitation onto those located in L3 following the CPP.

**Figure 2.**
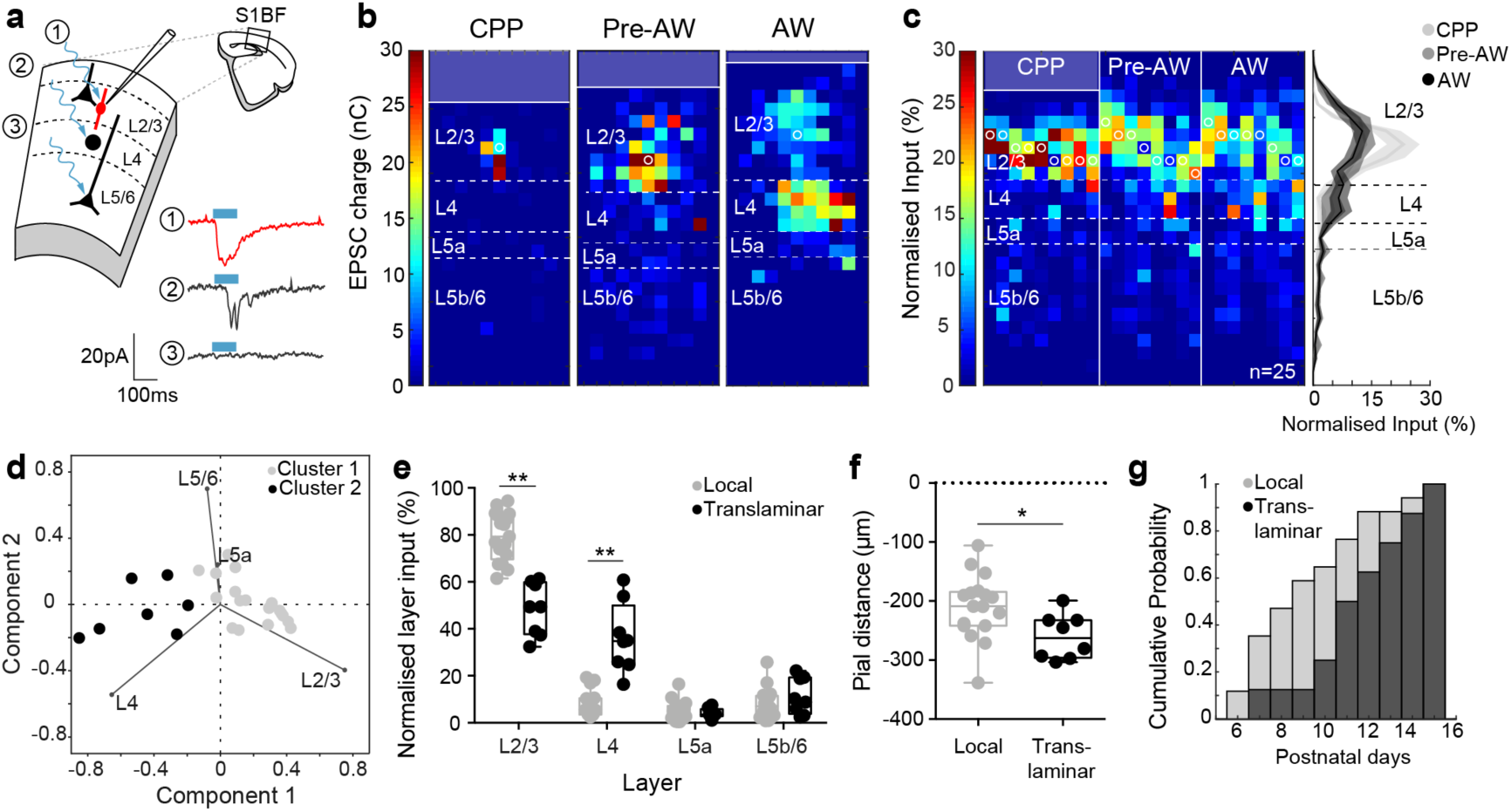
LSPS uncaging of glutamate reveals the emergence of distinct local and translaminar VIP+ IN populations following the L4 critical period for plasticity (CPP) (a) Schematic representation of laser-scanning photostimulation combined with whole-cell patch clamp, with current traces from different positions: (1) direct glutamate response time locked to laser onset; (2) synaptic response (EPSCs) at a delay to laser firing; (3) no LSPS-evoked response. Blue bar, 100ms UV laser pulse. (b) Glutamatergic afferent input maps for 3 VIP+ INs recorded across the different time windows tested. White circles indicate location of cell soma; dashed lines indicate layer boundaries. (c) Afferent input profiles for all VIP+ INs recorded through development (n = 25 cells); profiles were aligned by the L3-L4 border and ordered by age. (c, right) Average normalized laminar profiles of glutamatergic inputs onto L2/3 VIP+ INs across development. (d) Scatter plot of the first two principal components scores following principal component analysis (PCA) of normalized glutamatergic layer input of VIP+ INs. The lines indicate different variables inserted in the analysis (i.e. the relative layer component). Clusters are indicated by different colors. (e) Normalised laminar distribution of glutamatergic inputs onto L2/3 VIP+ INs grouped by clusters shows the presence of a local and a translaminar population of L2/3 VIP+ INs. Two-way RM ANOVA followed by Sidak’s multiple comparisons **:P<0.0001 (f) Distance from pial surface of VIP+ INs grouped by clusters shows that translaminar cells are significantly deeper than local VIP+ INs. Two-tailed Mann Whitney test, *:P<0.05 (g) Age analysis of VIP+ INs according to the clusters identified suggests that the translaminar cluster appears later in development.

### Long-range synaptic inputs from anterior-motor cortex onto layer 2 VIP+ INs are present within the first postnatal week

VIP+ INs are key integrators of long-range synaptic communication in the adult neocortex (Lee et al., 2013; Wall et al., 2016; Zhang et al., 2014). To assess the timeline over which S1BF VIP+ INs receive long-range inputs, we adapted a viral strategy (Arruda-Carvalho et al., 2017) to express channelrhodopsin-2(H134R)-YFP (ChR2) in anterior-motor pyramidal cells in early development (**Figure 3a**). We first confirmed that ChR2 expression was sufficient to evoke light-dependent suprathreshold responses at the onset of the CPP window in anterior-motor pyramidal cells (**Figure 3b**). In addition, we observed robust YFP expression in a distinctive pattern in S1BF with clear innervation of infragranular layers, L4 septa and L1 (**Figure 3c**) as well as dense arborization of YFP-labelled axons surrounding TdTomato+ VIP+ INs (**Figure 3d**). We recorded from both tdTomato+ VIP+ INs and non-TdTomato pyramidal cells (**Figure 3d, right**) across the depth of L2/3 in S1BF in acute *in vitro* coronal slices and tested for long-range glutamatergic synaptic input from anterior-motor areas using widefield blue (470nm) LED light pulses. S1BF neurons were identified as receiving anterior-motor synaptic input (connected) if we observed short latency EPSCs, time-locked to the LED pulse (**Figure 3e**). To exclude the possibility that an absence of such short latency EPSCs arose from a failure of our optogenetic strategy, neurons were only defined as unconnected if we observed either long latency, polysynaptic EPSCs in recorded cells or found connected cells in the same slice; both evidence of functional levels of ChR2 expression in afferent fibers in S1BF. We found that while both VIP+ INs (31%) and PYRs (62%) receive long-range inputs during CPP (**Figure 3f**), long-range connectivity onto VIP+ INs significantly increase by AW (82% of VIP+ INs), while remaining stable for PYRs. In contrast to adult cortex (Lee et al, 2013), we observed no difference in the amplitude of anterior-motor EPSCs onto VIP+ INs and PYRs during early development (**Figure 3g**). Moreover it was apparent that during CPP, anterior-motor synaptic input formed preferentially onto superficial, presumptive L2 neurons irrespective of cell type (**Figure 3h**). This further supports segregation in the function of early L2 and L3 VIP+ IN circuits, with L2 VIP+ INs integrating long-range signals while L3 VIP+ INs are progressively engaged by feed-forward excitation from the immediate column.

**Figure 3.**
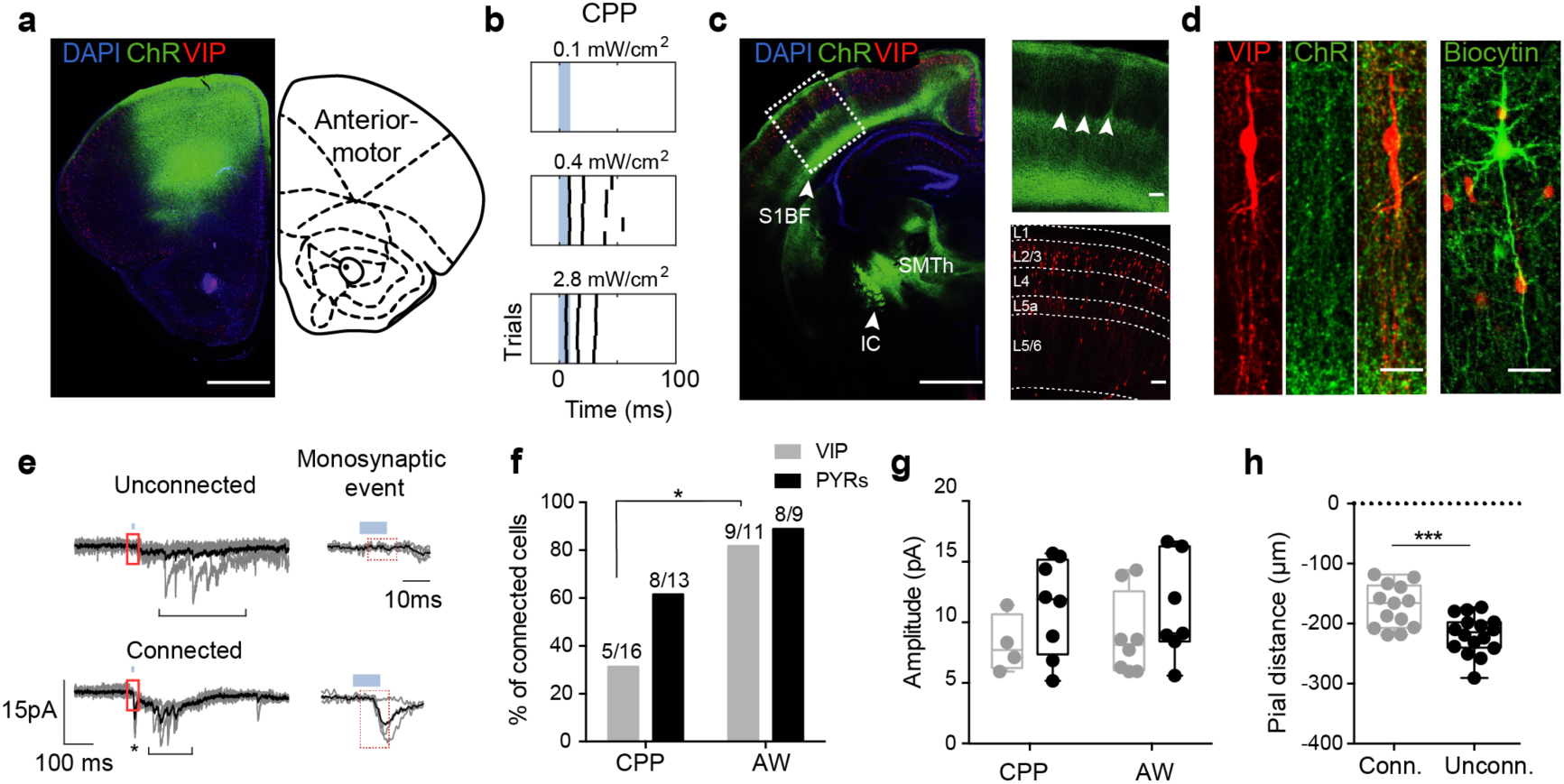
Long-range synaptic inputs from anterior-motor areas onto S1BF VIP+ INs as early as CPP. (a) Injection site during CPP, 7 days post injection; right panel adapted from Paxinos et al., (2007). Scale bar, 1mm. (b) Raster plot showing the timing of light-evoked action potentials in a ChR2+ cell (recorded in cell-attached mode) in anterior-motor areas at CPP at different LED powers. Blue bar, duration of LED pulse. (c) Recording site at CPP, 7 days post injection. Dashed box indicates area of S1BF shown at higher magnification in right panels. Arrowheads in higher magnification image indicate the fibers innervating septa between barrels. Scale bars, 1mm in the main picture and 100 µm in the higher-magnification pictures. Dashed lines: layer boundaries. S1BF, somatosensory barrel field; SMTh, sensory-motor cortex related thalamus; IC, internal capsule. (d, left) Image of a single VIP+ IN from panel (c) surrounded by ChR2-YFP fibers. Scale bar, 25µm. (d, right) Recovered morphology of a L2/3 pyramidal (PYR) cells. Scale bar, 25 µm. (e) Superimposed voltage clamped traces of unconnected and connected VIP+ INs at CPP following repeated wide-field 470nm LED stimulation. Blue bar, duration of LED pulse; asterisk, monosynaptic response; bracket, polysynaptic events. Area enclosed by the red box is shown magnified on the right: red dashed line, monosynaptic event time window. Grey, single sweep; Black, average response. (f) Proportions of S1BF VIP+ INs and pyramidal cells (PYRs) that receive monosynaptic connections from anterior-motor areas during CPP and AW time windows. Fisher’s exact test, * P ≤ 0.05 (g) Amplitude of minimal stimulation EPSCs in VIP+ INs and pyramidal cells. Two-way ANOVA, F (1, 23) = 0.1123, P>0.05 (h) Analysis of distance from pial surface for connected neuron (both VIP+ INs and PYRs) during CPP, two-tailed; unpaired t test *** P<0.001.

### VIP+ INs form synapses onto postsynaptic targets following an inside-out pattern during postnatal development

VIP+ INs are thought to influence cortical circuits primarily through disinhibition of PYRs via SST+ GABAergic INs, (Lee et al., 2013; Pfeffer et al., 2013; Pi et al., 2013; Staiger et al., 2004). To investigate the emergence of VIP+ IN output in S1BF we crossed our *VIP-Cre* line onto a background containing the ChR2-EYFP reporter allele (*Ai32*)(Madisen et al., 2012) and the *Lhx6-EGFP* BAC transgene that labels both multipolar, putative parvalbumin-expressing (PV+) and bitufted, putative-SST+ INs derived from the *Nkx2-1*-expressing medial ganglionic eminence (Gong et al., 2003) (**Figure 4a**). To confirm robust levels of ChR2 expression in VIP+ INs, sufficient for *in vitro* optogenetics, we performed extracellular recordings from EYFP+ cells and established that we could evoke action potentials upon blue (470nm) light illumination from CPP onward (**Figure 4b**). We then recorded from putative postsynaptic targets of VIP+ INs across the depth of neocortex, targeting PYRs and GABAergic INs in non-*Lhx6-EGFP and Lhx6-EGFP*+ offspring respectively. Postsynaptic neurons were voltage clamped at 0mV and the presence (connected) or absence (unconnected) of short latency, time locked inhibitory post-synaptic currents (IPSCs) from VIP+ INs established using brief pulses of wide-field blue light illumination (**Figure 4c**). We observed synaptic input from VIP+ INs onto both PYRs and *Lhx6-EGFP*+ INs across development (**Figure 4d**). The incidence of VIP+ input onto PYRs increased over development, but remained constant onto *Lhx6-EGFP*+ INs. However, we observed an inside-first outside-last pattern of innervation with an increased probability of connection onto infragranular as opposed to supragranular *Lhx6-EGFP*+ neurons during CPP (**Figure 4e**). Although, we cannot discount innervation by the small proportion of infragranular VIP+ INs onto local *Lhx6-EGFP*+ INs at this time, recovered morphologies of L2/3 VIP+ INs (**Figure 4f**) during CPP revealed prominent descending axons innervating infragranular layers from CPP onward. During AW time window, VIP+ INs showed increased connectivity onto L2/3 neurons irrespective of subtype, both in terms of connection probability (**Figure 4g**) and IPSC amplitude (**Figure 4h**). These observations suggest that VIP+ INs influence infragranular GABAergic circuits during the first postnatal week at a time point when interneurons form transient translaminar connections that regulate emergent thalamocortical networks (Marques-Smith et al., 2016).

**Figure 4.**
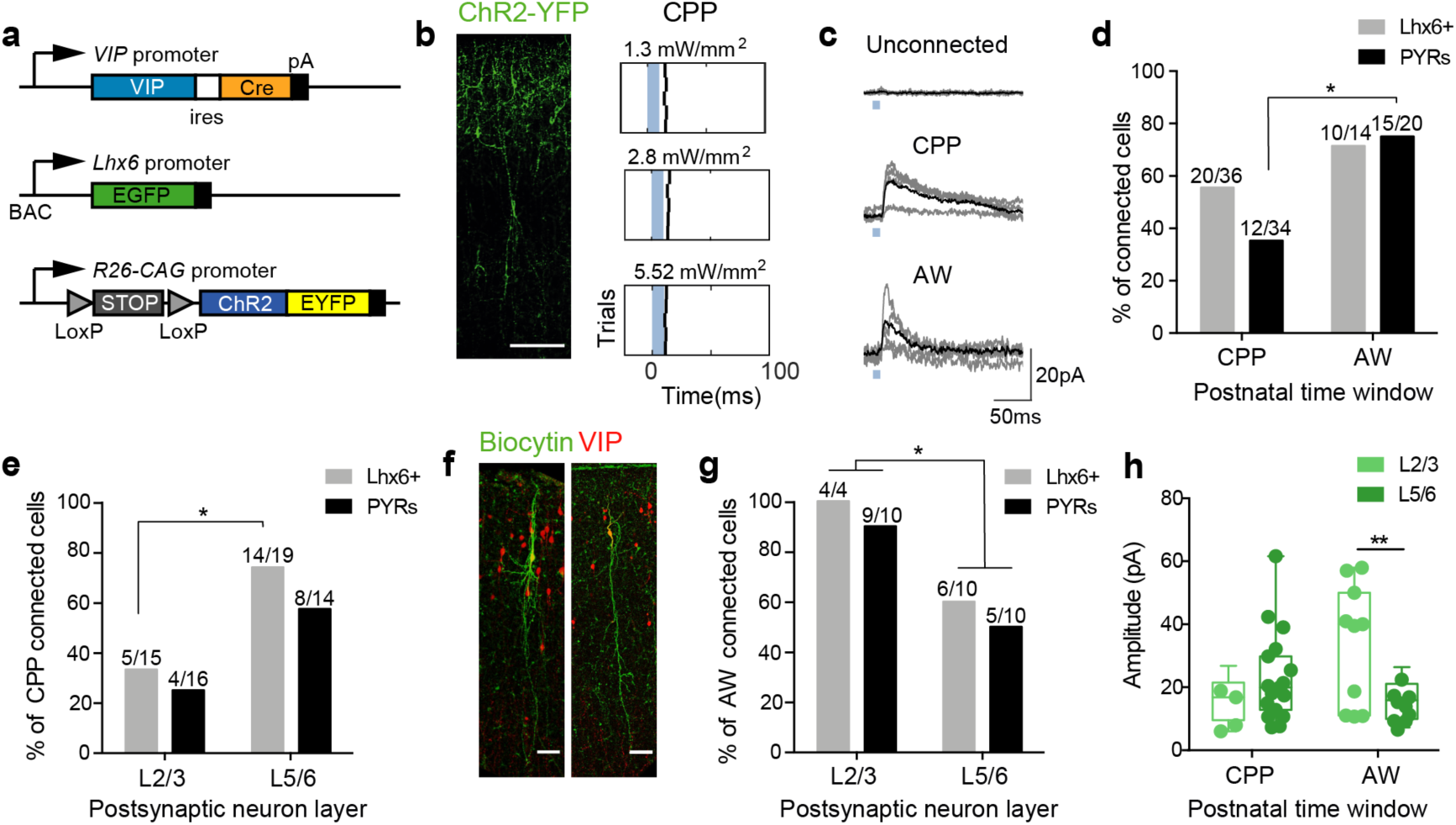
VIP+ INs form synaptic connections onto *Lhx6-EGFP* INs and PYRs through early postnatal development in S1BF. (a) Conditional genetic strategy to assess VIP+ IN connectivity over development; ires, internal ribosome entry site; pA, polyadenylation signal. R26-CAG, CMV early enhancer/chicken β actin promoter under the ROSA26 locus. (b) Immunohistochemistry (left) showing ChR-YFP+ cells in the *VIP-Cre;Ai32* line. Scale bar: 100µm. Raster plots (right) obtained from loose cell-attached recordings from a CPP VIP+ IN showing time-locked responses following stimulation at increasing LED power (n=5 trials under each condition). Blue bar, duration (10ms) of LED pulse. (c) Electrophysiological current traces showing unconnected (top trace) as well as connected *Lhx6-EGFP*+ cells receiving VIP+ IN synaptic input at CPP (middle) and AW (bottom). (d) Proportion of *Lhx6-EGFP*+ and PYRs recruited during the CPP and AW time windows irrespective of layer location. Fisher’s exact test with Bonferroni-Holm correction *P<0.05 (e) Layer distribution of VIP+ IN input onto *Lhx6-EGFP*+ INs and PYRs during CPP. Fisher’s exact test, *P<0.05. (f) Recovered morphologies of VIP+ INs recorded during CPP show axons descending into infragranular layers. (g) Proportion of *Lhx6-EGFP*+ and PYRs across layers receiving VIP+ IN synaptic input during the AW time window. Fisher’s exact test, *P<0.05. (h) Amplitude of ChR-evoked VIP+ IN IPSCs across layers, irrespective of cell type, during the CPP and AW time windows. Sidak’s multiple comparisons following two-way ANOVA **P<0.01.

### Cell-autonomous deletion of the transcription factor *Prox1* alters VIP+ IN synaptic integration into local but not long-range circuits

To further understand the contribution of VIP+ INs to emergent sensory processing we conditionally deleted the transcription factor *Prox1*, a regulator of postmitotic maturation in CGE-derived INs (Miyoshi et al., 2015), in a cell-autonomous fashion. Specifically, we wanted to examine if *Prox1* deletion impacted local and/or long-range synaptic integration of VIP+ INs and determine the consequences for VIP+ Ins function. We bred *Vip-Cre*^*HOMO*^;*Prox1*^*C/*+^ males with *Ai9*^*HOMO*^;*Prox1*^*C/C*^ females to generate offspring in which VIP+ INs were *Prox1* conditional knock-out (*Prox1*^*C/C*^; cKO) and labelled with tdTomato (**Figure 5a,b**). Due to the reported effects of *Prox1* haploinsufficiency in other systems (Harvey et al., 2005) we excluded *Prox1* heterozygous (*Prox1*^*C/*+^) animal from our final analysis. As such, we assessed the distribution of VIP+ INs in cKO animals compared to wild-type (WT, **Figure 5c,d**) and observed an increase in L5b VIP+ INs at P21 following *Prox1* cell-autonomous deletion, although the broad distribution was still biased towards supragranular layers. Analysis of VIP+ IN distribution across the depth of cortex further identified a difference in skewness between WT (skewed toward L2) and cKO animals, suggesting that in the latter VIP+ INs are more broadly distributed across the depth of the cortex (**Supplemental Figure 5a**) similar to previous reports of altered migration of VIP+ INs following *Prox1* deletion (Miyoshi et al., 2015). In addition, recovered morphologies of recorded VIP+ INs from cKO animals were consistent with previous findings (Miyoshi et al., 2015) in that cells did not display characteristic bipolar morphologies (**Figure 5e**).

**Figure 5.**
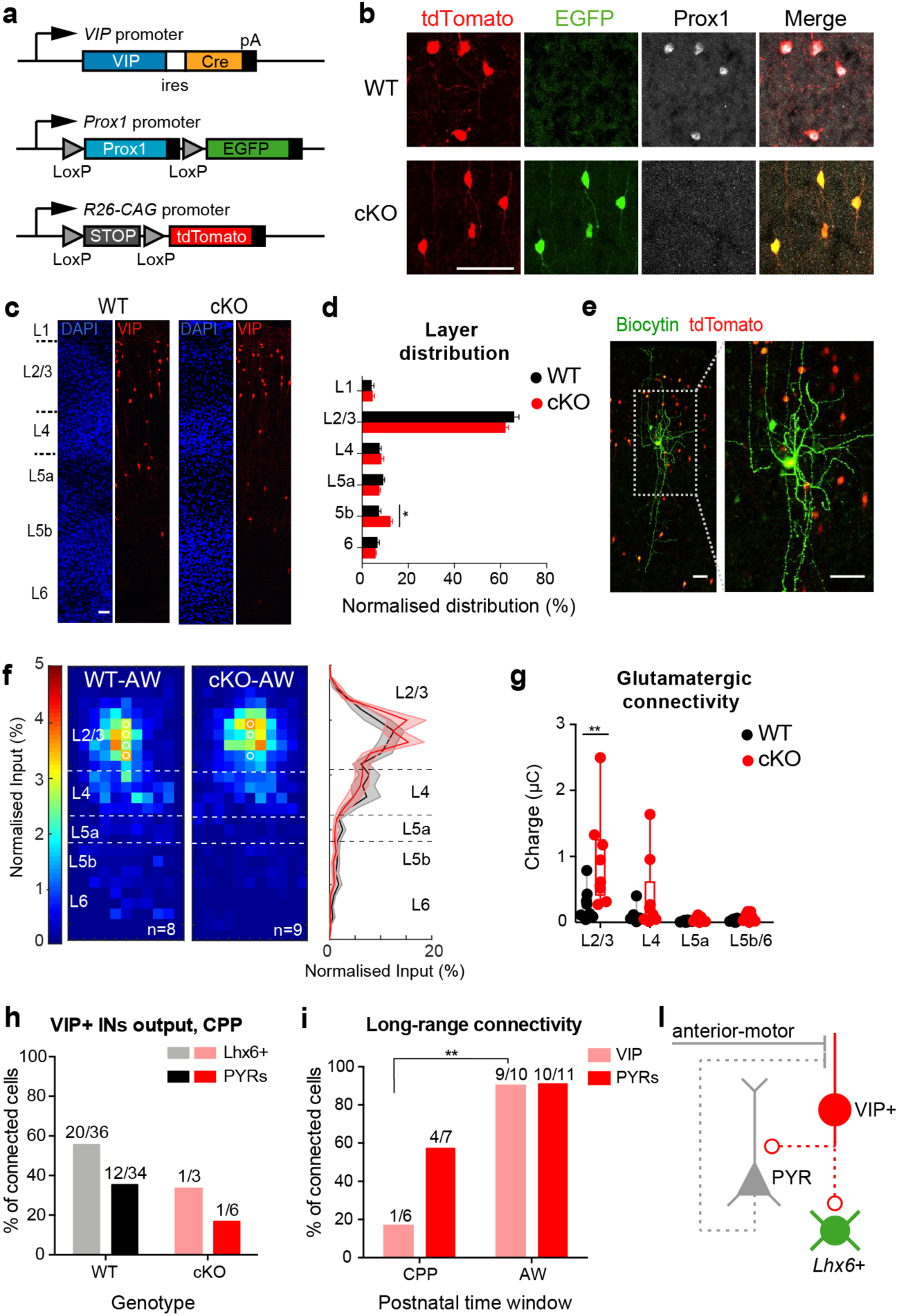
Genetic impairment of VIP+ INs leads to circuit defects *in vitro*. (a) Genetic strategy employed to conditionally delete *Prox1* in VIP+ INs. (b) Immunohistochemistry showing effective deletion of Prox1 and expression of EGFP in P21 cKO (*VIP-Cre;Ai9;Prox1*^*C/C*^) versus WT (*VIP-Cre;Ai9)*. Scale bar 50µm. (c) Location of VIP+ INs (red) across the depth of cortex in wild-type (WT) and cKO animals at P21. Scale bar 50µm. (d) Layer distribution of VIP+ INs. Sidak’s multiple comparisons test following two-way ANOVA *P<0.05. Control, 9 counts, n=3 animals; cKO, 8 counts n=3 animals. (e) VIP+ INs morphologies showed kinked processes following deletion of *Prox1* as previously reported (Miyoshi et al., 2015); scale bar 50µm (f) Average normalised glutamatergic synaptic input maps onto L2/3 VIP+ INs in WT and cKO animals at AW; maps aligned on the L3-L4 border. White circles depict the location of cell bodies; dashed white lines demarcate average layer boundary. (Right panel) average normalised laminar profiles of WT and cKO cells. (g) Layer distribution of glutamatergic input for WT and cKO L2/3 VIP+ INs; Two-way ANOVA followed by Sidak’s multiple comparisons test, **P<0.01 (h) Percentage of *Lhx6-EGFP*+ and PYR cells receiving VIP+ INs synaptic input during CPP in WT and cKO animals (i) Percentage of VIP+ INs receiving monosynaptic connections from anterior-motor areas in cKO animals. Fisher’s Exact Test **P<0.01 (l) Altered (dashed line) and normal (solid line) synaptic connections in the cKO VIP+ IN circuit. Flat line endings, glutamatergic synapses; circle endings; GABAergic synapses.

To examine the impact of *Prox1* deletion on local synaptic integration of VIP+ INs, we performed LSPS glutamate uncaging to determine PYR afferent input onto VIP+ INs in cKO versus WT. At CPP and pre-AW time points we observed mainly local glutamatergic inputs in cells recorded from cKO animals (**Supplementary Figure 5b**) similar to the majority of WT VIP+ cells at the same ages (**Figure 2**). However, during AW, the normalised input maps and layer input profile (**Figure 5f**) revealed a shift toward local L2/3 input in cKO animals. Analysis of average charge per layer confirmed an increase in local input from L2/3 PYRs onto VIP+ INs in cKO versus WT animals (**Figure 5g**). These data point to a disruption in the synaptic integration within the local network of cKO VIP+ INs in early development. To assess if this extended to efferent targets of VIP+ INs, we then crossed our conditional *Prox1* allele onto the *Ai32;Lhx6-EGFP* background and tested for the presence of ChR2-evoked IPSCs in both *Lhx6-EGFP*+ INs and PYRs. We found reduced connectivity onto both populations as early as CPP (**Figure 5h**) highlighting further deficits in local synaptic integration as a result of *Prox1* deletion. In contrast, long-range input from anterior-motor areas – assessed using viral transduction of anterior-motor pyramidal cells with ChR2 – revealed a similar timeline in emergent connectivity (**Figure 5i**) compared to WT (**Figure 3**) with VIP+ INs progressively recruited across the first two postnatal weeks. Taken together, these findings support previous observations that cell autonomous deletion of *Prox1* alters the synaptic integration of VIP+ INs (Miyoshi et al., 2015). They further identify subtle alterations in local connectivity within the immediate column without discernible impact on long-range connections (**Figure 5l**).

### Consequences of altered VIP+ IN synaptic integration on emergent S1BF function *in vivo*

VIP+ INs play important roles in sensory perception in S1BF (Barson et al., 2020; Lee et al., 2013; Sachidhanandam et al., 2016; Yu et al., 2019). To understand the long-lasting consequences of our genetic perturbation of VIP+ INs, we performed *in vivo* extracellular recordings in S1BF of anaesthetised juvenile (P21) animals to ascertain the impact of *Prox1* deletion on spontaneous activity (baseline) and multi-whisker evoked sensory activity. Placement of electrodes in S1BF was confirmed by short latency local field potential (LFP) deflections in response to multi-whisker stimulation and *post hoc* recovery of DiI track through the whisker barrel field (**Figure 6a**). We observed an increase in both the spontaneous multi-unit activity (MUA) across all layers of neocortex (**Figure 6b,c**) as well as in the power density of slow waves (0.5 – 2Hz) (**Figure 6d**) in cKO as opposed to WT animals.

**Figure 6.**
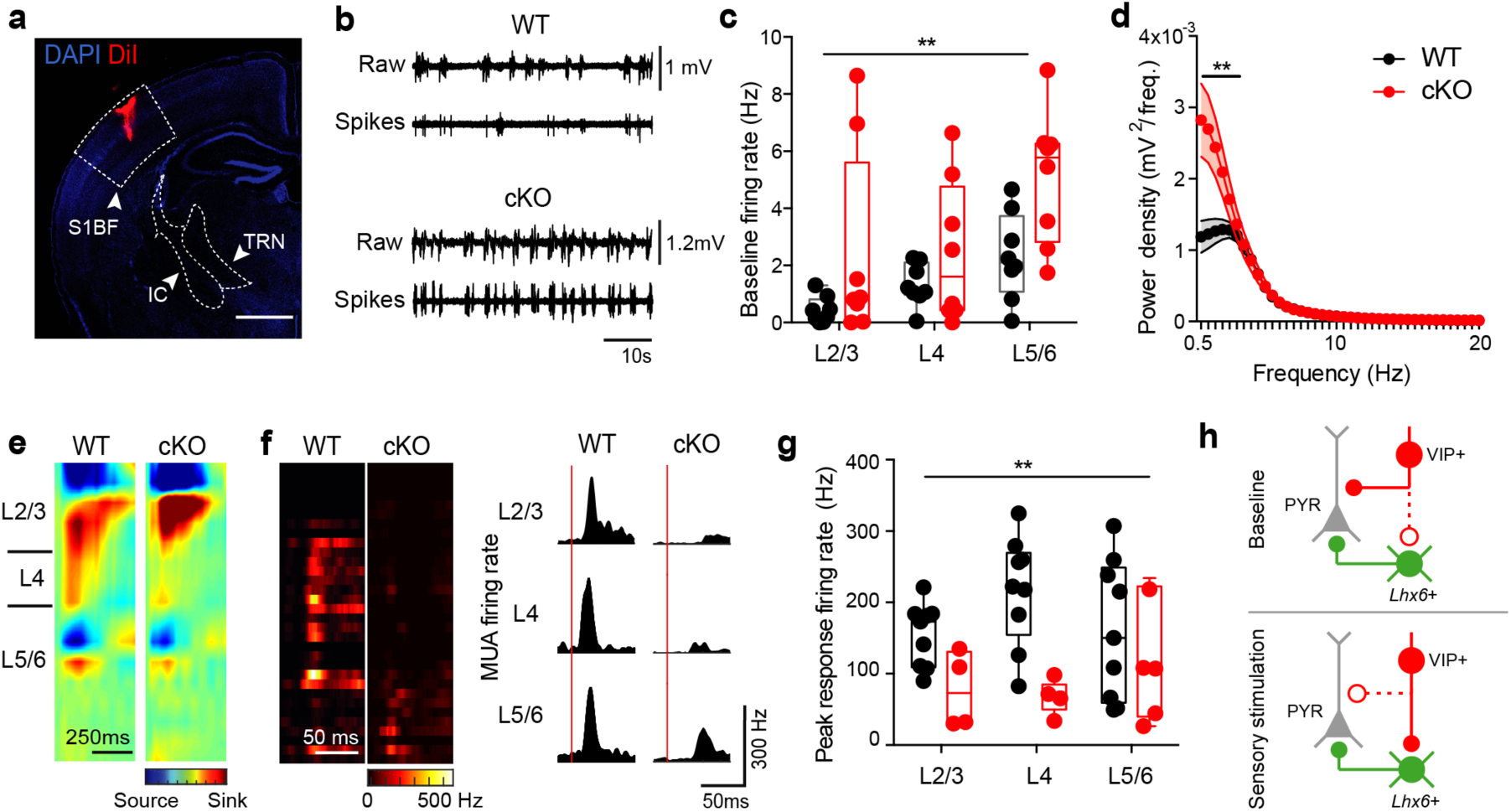
Conditional deletion of *Prox1* in VIP+ INs leads to altered network activity *in vivo*. (a) *Post hoc* histological analysis to confirm electrode position following extracellular recording; scale bar 1mm. TRN, thalamic reticular nucleus; IC, internal capsule. (b) Local field potential (LFP) traces and multi-unit activity (MUA) during baseline activity in WT and cKO animals. (c) Average spontaneous MUA frequency across the depth of the cortex in WT and cKO animals; Two-way ANOVA, F (1, 42) = 10.62, **P<0.01 (d) LFP power analysis in WT and cKO reveals genotype-dependent differences across the 0.5 – 2Hz frequency bands; Two-way RM ANOVA, F (39, 702) = 6.895, **P<0.0001 (e) Current-source density (CSD) maps for WT and cKO across layers following whisker stimulation. The CSD plot is visualized as a pseudo-color image with red representing sinks and blue indicating sources. (f) (left) MUA heat maps following multi-whisker stimulation for WT and cKO. (Right) layer-specific multi unit activity following multi-whisker stimulation (onset indicated by the red bar). (g) Average peak multi-unit activity (MUA) upon multi-whisker stimulation across the depth of the cortex in WT and cKO animals; Two-way ANOVA, F (1, 36) = 13.99, **P<0.001. (h) Model of the action of VIP+ INs in anaesthetised animals during baseline and sensory stimulation. Solid line, active synaptic pathway; dashed line, quiescent connection.

Multi-whisker stimulation in both WT and cKO animals resulted in characteristic sink-source pattern of recruitment across the layers of S1BF (**Figure 6e**) and prominent short latency MUA in L4 and L5/6 of WT animals (**Figure 6f**). In contrast to the increased spontaneous activity in cKO mice, whisker stimulation resulted in a decrease in peak MUA firing rate in cKO as opposed to WT animals (**Figure 6f**) with an overall effect observed across all layers due to genotype (**Figure 6g**). We conclude that deletion of *Prox1* in VIP+ INs results in contrasting effects on spontaneous and sensory-evoked *in vivo* activity (**Figure 6h**).

## DISCUSSION

We have used optical strategies in combination with electrophysiology to define postnatal synaptic integration and function of VIP+ INs in mouse somatosensory cortex. We found that VIP+ INs integrate into local and anterior-motor circuits as early as the L4 critical period of plasticity (P5-8). These data identify an early role for this interneuron population that suggests that they contribute to early cortical circuit maturation and plasticity. Perturbation of the genetic specification of VIP+ INs through conditional deletion of the transcription factor *Prox1* impairs local circuit integration and somatosensory processing.

VIP+ INs are a major class of GABAergic interneuron that originate from the caudal ganglionic eminence (CGE). CGE INs, defined by expression of the ionotropic 5-HT_3A_ receptor as well as the transcription factor *Prox1* (Lee et al., 2010; Miyoshi et al., 2015; Rubin and Kessaris, 2013), have a late birthdate compared to the other two major classes of GABAergic IN – PV+ and SST+ INs – that originate from the medial ganglionic eminence (MGE) (Butt et al., 2005; Miyoshi and Fishell, 2011; Miyoshi et al., 2010). As such, CGE INs are regarded as late integrators into the cortical circuitry that preferentially populate supragranular layers of neocortex, modulate other INs – notably SST+ cells – and play a role in mediating top down influences (Fu et al., 2014; Karnani et al., 2016; Lee et al., 2013; Pfeffer et al., 2013; Pi et al., 2013; Prönneke et al., 2015; Wall et al., 2016; Zhang et al., 2014). However our data identify a more complex picture, with VIP+ INs influencing infragranular MGE-derived INs and pyramidal cells within the first postnatal week in S1BF; a time point when VIP+ INs have not fully integrated into either the local, columnar or long-range glutamatergic network. Following the layer 4 critical period for plasticity (CPP; P5-P8), we observe the emergence of translaminar glutamatergic connections on supragranular VIP+ INs as well as concurrent increase in synaptic input from anterior-motor areas, such that the mature VIP+ IN circuit is largely in place by the onset of active perception and exploratory behaviour around P12 (Arakawa and Erzurumlu, 2015). Our data suggest that a distinction can be made between L2 and L3 VIP+ INs at early ages, with the latter receiving emergent feed-forward excitation from L4 and the former recipient of anterior-motor input at early ages. Cell autonomous deletion of the transcription factor *Prox1* in VIP+ INs results in altered migration and synaptic integration into the immediate column but spares long-range connectivity. These impairments at the network level underlie an increase in spontaneous *in vivo* network activity with a parallel increase in slow-wave power density band (0.5-2Hz) present under urethane anaesthesia (Clement et al., 2008). In contrast, whisker-evoked activity is attenuated across the depth of cortex that could arise from reduced VIP+ IN disinhibition of pyramidal cell following conditional deletion of *Prox1*. This broad effect across the layers of neocortex is at odds with previous work, that has identified layer-specific VIP+ IN disinhibition in S1BF (Muñoz et al., 2017) but further supports the idea that *Prox1* is necessary to establish such layer-specific VIP+ IN networks (Miyoshi et al., 2015). In addition our *in vivo* observations match those of others which have reported differing effects on spontaneous and sensory evoked activity (Batista-Brito et al., 2017; Mossner et al., 2020).

Coordinated activity between motor and sensory areas starts early in development (An et al., 2014; Luhmann, 2017; McVea et al., 2012, 2017), prior to the onset of active whisking at the end of the second postnatal week (Arakawa and Erzurumlu, 2015; Bureau et al., 2004). We have used a viral optogenetic strategy, previously employed to delineate the emergence of prefrontal cortex-amygdala connectivity (Arruda-Carvalho et al., 2017), to show that long-range connections from anterior-motor areas engage not only with pyramidal cells in S1BF, but also VIP+ INs (Lee et al., 2013; Wall et al., 2016) as early as the CPP. This extends our knowledge of how brain-wide circuits come online (Arruda-Carvalho et al., 2017; De León Reyes et al., 2019) and identifies the recruitment of GABAergic interneurons during circuit plasticity and maturation across brain areas. The relative balance between long-range and local influences remains to be investigated but our data argues for an initial compartmentalization of these two influences prior to the potentiation of feed-forward and local synapses in supragranular layers at the onset of active perception (Bureau et al., 2004; Clem and Barth, 2006; Clem et al., 2008; Itami and Kimura, 2012; Wen and Barth, 2011).

To assess the efferent targets of VIP+ INs through development we used a conditional optogenetic strategy and recorded from candidate postsynaptic neurons (Garcia-Junco-Clemente et al., 2017; Lee et al., 2013; Pi et al., 2013) including MGE-derived INs identified using the Lhx6-EGFP transgenic line. We observed that VIP+ INs form functional synapses onto postsynaptic neurons in an inside-out manner and target infragranular pyramidal cells and Lhx6-EGFP INs from the CPP onward – including SST+ interneurons that we and others have shown to be important for circuit maturation (Marques-Smith et al., 2016; Tuncdemir et al., 2016). Indeed, it is evident that the pattern of VIP+ IN innervation of post-synaptic targets shifts with the progressive inside-out maturation of the cortical circuitry in line with the emergence of the canonical cortical circuit (Bureau et al., 2004; Erzurumlu and Gaspar, 2012; Marques-Smith et al., 2016; Wen and Barth, 2011). Such that during the latest time window we studied, AW, VIP+ IN innervation of postsynaptic neurons matches the pattern observed in adult cortex (Muñoz et al., 2017; Zhou et al., 2017; Feldmeyer et al., 2018). While we cannot discriminate the specific subtype of the postsynaptic MGE interneurons that receive VIP+ IN input, these likely include both PV+ and SST+ INs as previously reported (Dávid et al., 2007; Hioki et al., 2013; Lee et al., 2013; Pfeffer et al., 2013; Pi et al., 2013; Staiger et al., 2004; Zhang et al., 2014), and early SST+ IN networks important for circuit maturation (Marques-Smith et al., 2016; Tuncdemir et al., 2016). This further suggests that VIP+ INs are capable of disinhibition – a proven mechanism for synaptic plasticity (Fu et al., 2015; Letzkus et al., 2015; Pi et al., 2013; Williams and Holtmaat, 2019) from the CPP onward. Moreover our mapping of the efferent targets of VIP+ IN reinforce the body of evidence showing that this class of neuron target pyramidal cells directly (Chiu et al., 2018; Garcia-Junco-Clemente et al., 2017; Lee et al., 2013; Pfeffer et al., 2013) and do so from an early age. As such VIP+ INs are, from the moment that they integrate into neocortex, well positioned to exert varied effects as described in the juvenile and adult cortex (Batista-Brito et al., 2017; Bigelow et al., 2019; Cardin, 2019; Dipoppa et al., 2018; Fu et al., 2014; Garrett et al., 2020; Keller et al., 2020; Millman et al., 2019; Pakan et al., 2016; Pi et al., 2013).

We tested the requirement for VIP INs using a genetic strategy – conditional loss of function of the transcription factor *Prox1*, that has previously been shown to alter the migration and synaptic integration of CGE-derived, 5-HT_3A_R+ INs (Miyoshi et al., 2015). Our results point to subtle deficits in the layer location and synaptic integration of VIP+ INs within the local S1BF circuit following cell autonomous deletion of *Prox1* in VIP+ INs; deficits that have consequences for sensory processing in the juvenile mouse. The selective impact on local connections points to a failure of VIP+ INs to appropriately interpret normal columnar signals for synaptic integration (Miyoshi et al., 2015; Wester et al., 2019) following deletion of this transcription factor. Altered molecular machinery could include members of the neurexin – neuroligin synaptic organizer protein family that are selectively expressed by *Prox1*+ INs (Lukacsovich et al., 2019), and have the ability to differentially regulate local versus long-range glutamatergic connections (Pregno et al., 2013). That aside, deletion of *Prox1* in VIP INs results in attenuation of multi-whisker evoked activity across the depth of cortex, pointing to impairment in the processing of incoming sensory information (Barson et al., 2020; Lee et al., 2013; Sachidhanandam et al., 2016; Yu et al., 2019).

Taken together, our observations align with a number of recent reports suggesting that genetic perturbation of VIP+ INs early in development can have significant effects on cortical processing (Batista-Brito et al., 2017; Goff and Goldberg, 2019; Mossner et al., 2020; Qiu et al., 2020). We propose that VIP+ INs represent a core component of early GABAergic networks in S1BF, one that is able to integrate sensory and motor information within the first postnatal week. We identify that through these synaptic connections, and interactions with ascending pathways including thalamic input (Che et al., 2018) and neuromodulatory systems (De Marco García et al., 2015; Frazer et al., 2015; Lee et al., 2010; Murthy et al., 2014), VIP+ INs are well positioned to influence the plasticity and maturation of both GABAergic and glutamatergic cortical networks from the first postnatal week onward.

## METHODS

### Animal husbandry and use

The following mouse lines were maintained on a mixed C57BL/6J and CD1 background: *VIP-ires-Cre* (*Vip*^*tm1(cre)Zjh*/^J) (Taniguchi et al., 2011); *Lhx6-EGFP* (Tg(Lhx6-EGFP)BP221Gsat) (Gong et al., 2003); *Ai9* (B6.Cg-*Gt(ROSA)26Sor*^*tm9(CAG-*^ *tdTomato)Hze*/J) (The Jackson Laboratory); *Ai32* (B6;129S-*Gt(ROSA)26Sor*^*tm32(CAG-*^ *COP4*H134R/EYFP)Hze*/J) (The Jackson Laboratory); *Prox1* (Prox1^tm1.1Fuma^) (Iwano et al., 2012). Conditional *Prox1* mice generated by Prof. Fumio Matsuzaki (RIKEN Centre for Developmental Biology, Minatojima Minamimachi Chuo-ku Kobe Japan) were provided by Prof. Paul Riley (Oxford, UK). Animals were kept with the dam until weaning on a 12 h light/dark cycle with food and water provided *ad libitum*. Experimental animals were heterozygous for *VIP-Cre* and the reporter (either *Ai32* or *Ai9*), and, when applicable, hemizygous for *Lhx6-EGFP*. Conditional *Prox1* animals were homozygous for the *Prox1* floxed allele. *VIP-Cre;Ai9, VIP-Cre;Ai32;Lhx6-EGFP, VIP-Cre;Ai9;Prox1*^*C/C*^ neonates were obtained from appropriate breeding pairs and used for experiments between postnatal day 0 and 22 (P0 and P22). Animal use was approved by the local ethical review panel and conducted according to the Home Office project (30/3052 and P861F9BB75) and personal licenses under the UK Animals (Scientific Procedure) 1986 Act.

### Genotyping

All experiments involving the *Prox1* line (Prox1^tm1.1Fuma^) (Iwano et al., 2012) were performed blind to the genotype, which was confirmed by PCR following completion of data analysis. The following primers were used: Forward: CAG CCC TTT TGT TCT GTT GGC CAG; Reverse: GCA GAT GCT GTC CCT ACC GTC C. PCR was performed as follows: after 2 min of denaturation at 94 °C, 35 amplification cycles were performed (94 °C for 30 s, 60 °C for 30 s, 72 °C for 45 s), followed by a final extension stage at 72 °C for 5min. PCR products were kept at 4°C and then run on a 1.5% agarose gel in TAE (Tris-acetate-EDTA) buffer (Sigma, UK) with the conditional allele showing a larger band (220bp) versus wild-type (WT) (184bp)(Iwano et al., 2012).

### Immunohistochemistry

Following anaesthesia induction with 4% isoflurane in 100% O2, pups of both sexes were euthanized with an overdose (>200mg/kg) of 20% pentobarbital sodium solution and transcardially perfused with 4% paraformaldehyde (PFA, Alfa Aesar) in phosphate-buffered solution (PBS, Sigma). Brains were dissected out, post-fixed in 4% PFA in PBS for 1 hour at 4°C, washed in PBS and stored in PBS-Azide (0.02-0.05%) at 4°C until use. Brains were then prepared for vibratome or cryostat cutting. In the case of the former (e.g. tdTomato, GFP, and calretinin staining), brains were embedded in a 5% low-gelling agarose gel (in PBS) and 50µm coronal slices were cut using a vibroslicer (Leica VT1000S). Slices were then stored in PBS-Azide (0.02-0.05%) at 4°C. Alternatively (e.g. VIP and Prox1 staining), brains were cryoprotected using increasing concentrations of sucrose (10% and then 30%) in PBS at 4°C overnight. Brains were then embedded in O.C.T. compound (VWR) on dry ice and kept at −80°C until they were sectioned on a cryostat (Leica Biosystems) into 20 – 25µm serial coronal sections and mounted on SuperFrost Plus slides (VWR). Before immunohistochemistry, slices were washed in PBS at room temperature (RT) then permeabilized with PBS-T (0.1M PBS + 0.2% Triton-X100, Sigma) for 30 mins. prior to blocking with 5% normal goat serum (NGS, Invitrogen) in PBS-T for 1-2 hrs. at RT, and incubation with the primary antibody (polyclonal rabbit anti-dsRed antibody, 632496, Clontech, dilution 1:500; monoclonal mouse anti-dsRed antibody, 3994-100, BioVision, dilution 1:500; polyclonal chicken anti-GFP, ab13970, Abcam, dilution 1:2000; mouse monoclonal anti-Calretinin clone 6B8.2, MAB1568, Sigma-Aldrich, dilution 1:100; polyclonal rabbit anti-VIP, ab8556, Abcam, dilution 1:2000) in the blocking solution overnight at 4°C. For VIP staining, the primary antibody was kept for 2 nights at 4°C. After being washed in PBS and left additional 30 minutes in the blocking solution at RT, slices were incubated with a 1:500-1000 dilution of secondary antibody (goat anti-rabbit IgG (H+L) Alexa546 conjugate, A-11035, Molecular Probes; goat anti-chicken Alexa488 conjugate, ab150169, Abcam; goat anti-mouse IgG (H+L) Alexa568 conjugate, ab175473, Abcam; goat anti-rabbit Alexa633, A-21070, Molecular Probes) for 2 hours at RT. After 3×5 min washes in PBS, sections were counterstained with 4’,6-diamidine-2’-phenylindole dihydrochloride (DAPI, D3571, Molecular Probes, dilution 1:1000), re-washed, mounted when appropriate and cover-slipped with Fluoromount aqueous mounting medium (Sigma). Finally, they were permanently sealed using nail polish.

For *Prox1* staining, slices were permeabilised with PBS-T (0.1M PBS + 0.1% Triton-X100, Sigma) for 30 minutes and blocked with 10% normal goat serum (NGS, Invitrogen) in PBS-T for 1-2 hours at RT. Tissue was then incubated with the primary antibody (polyclonal rabbit anti-PROX1, AB5475, Sigma-Aldrich, dilution 1:250) in blocking solution overnight at 4°C. Following washing, the slices were incubated for 2 hours at RT with anti-rabbit biotinylated antibody (1:200) in blocking solution. After 3×10 minute washings, slices were incubated for 2 hours at RT with streptavidin Alexa488 conjugate (S11223, Molecular Probes, dilution 1:500). Washings, DAPI counterstaining and mounting were performed as previously described.

### Image acquisition and analysis

Microscopy imaging for cell counting was performed with a Zeiss laser scanning confocal microscope (LSM710), using a UPLS APO (air) 10x/0.40 objective at a pixel resolution of 1024 x 1024. Anatomically matched sections of the somatosensory barrel cortex were selected for cell counting, using the Atlas of the Developing Mouse Brain (Paxinos et al., 2007). For each image, cells were counted across all layers in regions of interest (ROI) of fixed width (either 500 or 1000 µm).

Imaging of Prox1 staining was obtained using an Olympus point-scanning confocal (FV3000), equipped with a UPLS APO (air) 20x/0.75 objective, at a pixel resolution of 1024 x 1024. For quantification of tdTomato+ cells, a customised macro developed in Fiji (Schindelin et al., 2012) was used. Briefly, the boundaries of the different cortical layers were determined by DAPI counterstaining. TdTomato signal was then used to automatically build a binary mask identifying only somata. To normalize the distribution of positive cells, each selected column was divided into 10 equal bins. Based on their centroid location, somata were then assigned to layers and bins. To quantify cell density, a similar strategy to the one reported by Prönneke et al. (2015) was followed. The area of the cortical depth in which cells were counted was measured (µm^2^) and multiplied by the thickness of the section, to obtain a volume value in µm^3^ which was then converted to a mm^3^ value.

### Viral injections

To ensure early channelrhodopsin-2 expression, the AAV1.CaMKIIa.hChR2(H134R)-eYFP.WPRE.hGH virus (Penn Vector Core, Addgene 269696P, Lot CS1322, titre 3.388 e^13^ GC/ml, suspended in PBS with 5% glycerol (Arruda-Carvalho et al., 2017)) was used. P0-P1 pups were separated from the dam, anesthetized through hypothermia for 5-8 min, and moved to a customised stereotaxic set-up. A small volume of virus (49.9nL x 3 cycles; rate: 0.23 nL/s; delay: 3 s) was injected with the Nanoject III (Programmable Nanoliter Injector, Drummond) using a pulled glass pipette (∼ 20 µm inner tip diameter, bevelled at a 45° angle). To ensure targeting of the developing anterior-motor areas, the following coordinates were used (calculated from vascular lambda and expressed as mm): AP −3.12; ML 1.1; DV 0.8. No incisions were made, but once reached the right AP and ML positions, the pipette tip was used to pierce through the skull and then positioned in the correct DV coordinates to perform the injection. The pipette was withdrawn ∼10-30 seconds after completion of injection. After 6-7 days post-injection (dpi) it was possible to observe good viral transduction of cell bodies in anterior-motor areas, as well as fibers present in S1BF. To confirm targeting of anterior-motor areas, perfused brains were sectioned into 50µm coronal slices on a vibrating microtome (Leica, VT1000S). Immunohistochemistry was performed using polyclonal chicken anti-GFP antibody (ab13970, Abcam, dilution 1:2000) and polyclonal rabbit anti-dsRed (632496, Clontech, dilution 1:500). Fluorescent images were acquired with a Zeiss 800 Airyscan using a 10x 0.45NA Plan-APOCHROMAT (Zeiss) objective.

### *In vitro* electrophysiology

Acute in vitro brain slices was prepared as previously described (Anastasiades et al., 2016; Marques-Smith et al., 2016). Mice of both sexes were used. Slices containing the somatosensory whisker barrel cortex (S1BF) were selected for electrophysiology experiments. Neurons ∼50µm below the slice surface were targeted for whole-cell patch-clamp recordings at RT using a MultiClamp 700B amplifier and a Digidata 1440A digitiser (Molecular Devices, USA). A standard potassium-based intracellular electrode solution of the following composition (in mM): 128 K-gluconate, 4 NaCl, 0.3 Li-GTP, 5 Mg-ATP, 0. 1 CaCl_2_, 10 HEPES and 1 glucose (pH 7.2 with KOH; 270-280 mOsm) was used to obtain intrinsic electrophysiological properties, as well as mapping of local and long-range glutamatergic inputs. To record GABAergic IPSCs, electrodes were filled with a Cesium-based intracellular solution, containing (in mM): 100 gluconic acid, 0.2 EGTA, 5 MgCl_2_, 40 HEPES, 2 Mg-ATP, 0.3 Li-GTP (pH 7.2 using CsOH; 270-280 mOsm). Intracellular solutions contained biocytin (3%) (Sigma UK) to enable recovery of the morphology of recorded cells. Glutamatergic EPSCs were recorded with the neuron voltage clamped at −60mV (Anastasiades and Butt, 2012). GABAergic IPSCs were recorded in cells voltage clamped at the reversal potential for glutamate (Eglut), set to 0 mV (corrected for calculated liquid junction potential) (Anastasiades and Butt, 2012; Anastasiades et al., 2016). Cortical layers were distinguished in the DIC image based on changes in cell size and density. Cell input and series resistance were monitored during the recordings without applying compensation; recordings were discarded when series resistance exceeded 20% of its initial value. Intrinsic electrophysiological properties were recorded in current clamp mode in response to depolarising or hyperpolarising current pulses of 500ms delivered at 0.5Hz via Clampex (v.10.1, Molecular Devices). Data was analyzed offline using a Matlab custom-written script.

### Laser scanning photostimulation (LSPS)

LSPS protocols were performed as previously described (Anastasiades and Butt, 2012; Anastasiades et al., 2016; Marques-Smith et al., 2016). Stimulation was performed using a low intensity (max. 2mW), long duration (100ms) laser pulse generated by an ultraviolet (UV) laser (DPSL-355/30) together with a UGA-42 targeting module (Rapp OptoElectronic GmbH) and focused through Zeiss Axioskop FS2-plus microscope equipped with a 10X UPLFLN objective (Olympus). Laser power was calibrated to the developmental age to ensure spatial resolution ∼50µm (Anastasiades and Butt, 2012; Anastasiades et al., 2016; Marques-Smith et al., 2016). Prior to LSPS, slices were incubated for at least 6 mins. with high divalent cation (HDC) ACSF (ACSF with 4mM MgCl_2_ and 4mM CaCl_2_, to reduce all polysynaptic and spontaneous activity), containing 100µM of 4-Methoxy-7-nitroindolinyl-caged-L-glutamate (MNI glutamate, Tocris Bioscience).

Photostimulation was triggered in a pseudorandom pattern to prevent sequential stimulation of adjacent sites at 1Hz (Anastasiades and Butt, 2012; Shepherd et al., 2003) and the position of the target grid adjusted to cover the entire depth of the cortex. LSPS grids at each locations were run multiple times, aiming to obtain a minimum of two-three repeat runs per cell. An IR-DIC photomicrograph of the LSPS grid relative to the patched cell was taken and used as a reference to reconstruct the pixel positions relative to the layer boundaries. Offline analysis of the current traces was performed adapting a customised Matlab script developed in the lab (Matlab2018a). Previously published criteria (Anastasiades and Butt, 2012) were used to determine the putative monosynaptic event detection window. The total charge was extracted for events captured within the monosynaptic window. Events with onsets before the monosynaptic window were considered to contain a direct-response component. In this case, the charge value was interpolated from all the other surrounding points using the Matlab function ‘griddata’ with linear interpolation (Weiler et al., 2018). This reduced overestimation of local connectivity caused by partially including the direct response. For each cell, the absolute (pC/pixel) and normalised (%pC/pixel) afferent input was measured. In normalised maps, each pixel indicates the amplitude of the average evoked EPSCs expressed as a percentage of the overall input evoked across the extent of the LSPS grid. Laminar input profiles were evaluated by summing each normalized input evoked from each horizontal row. Average maps were plotted aligning individual maps to the L3-L4 border.

### Cluster analysis of LSPS data

Principal component analysis (PCA) was performed on the normalised afferent input per layer obtained for each cell. Following identification of the first two principal components, cluster analysis was performed employing k-means clustering, using squared Euclidean distance as the metric and a maximum number of 1000 iterations. To determine the optimal number of clusters, silhouette analysis was performed.

### Optogenetic stimulation of VIP+ INs output

Activation of VIP+ INs in the *VIP-Cre;Ai32* line was achieved by wide-field blue light stimulation (Cool-LED pE-100, 470nm) focused through a 40x objective (Olympus LUMPLFLN 40XW Objective). 10ms square light pulses were delivered at multiple LED power intensity (0.082, 0.38, 0.7, 1.33, 2.83 and 5.52 mW/cm^2^). In each recording, stimulations were repeated five times every 20 seconds. Post-synaptic cells were considered responsive only if light-evoked, time-locked IPSCs were present in at least 2 out of 5 trials. The minimal stimulation, defined as the minimal LED power able to elicit at least 2 IPSCs in 5 trials, was calculated for each cell. A customised Matlab script (Matlab2018a) was used to extract percentage of failure and average response amplitude.

### Optogenetic stimulation of anterior-motor afferents in S1BF

Activation of virally expressed channelrhodopsin either in the soma of anterior-motor pyramidal cells or in their terminals in S1BF was achieved as outlined above. In this case, 1ms and 10 ms blue light pulses were delivered at different LED intensities to ensure to find the minimal stimulation required to engage postsynaptic cells. To confirm the monosynaptic nature of the responses and calculate the monosynaptic window, tetrodoxin (TTX, 1 µM) and 4-aminopyridine (4-AP, 100 µM) were added to the bath solution in a subset of cells (Petreanu et al., 2009; Suter and Shepherd, 2015; Yamawaki and Shepherd, 2015). Whenever EPSCs were not detected in L2/3 neurons, the presence of functional ChR2+ terminals was confirmed by recording a L6 pyramidal cell, an internal positive control as demonstrated by the abundant presence of ChR2+ fibers in L6. Data analysis was performed as described above.

### *In vivo* electrophysiology

Mice of both sexes aged P19-P22 were anaesthetised with 10% urethane in PBS (dose: 1 – 1.5g/kg) (Sigma). Mice were mounted in a stereotaxic frame (Kopf, with Stoelting pup adaptor) and the scalp removed to expose the skull. The following coordinates were used for S1BF, with lambda used as reference: AP 3.30 mm, ML 3.125 mm, DV −750 µm. A craniotomy of ∼2mm in diameter was obtained with a drill (Volvere i7, NSK Gx35EM-B OBJ30013 and NSK VR-RB OBJ10007) equipped with a 0.5 mm tip. A 32-channel single shank electrode (A1×32-Poly2-5mm-50s-177, Neuronexus) was slowly lowered into the brain. To ensure *post-hoc* confirmation of the electrode location, it was submerged in DiI (DiIC18(3), 2.5 mg/ml, D282 Lot 1990320 in 70% ethanol, Sigma) for 10-20 minutes before insertion. During the entire duration of the experiments, depth of anaesthesia and breathing were constantly monitored. The animal temperature was maintained with a heating pad (Watlow 025037500,120 Volts, 46 Watts, 1JR 1529C-14) set at 37 °C.

A 20 min baseline was recorded a few minutes after electrode insertion (Open Ephys acquisition board). A piezo stimulator (PB4NB2W Piezoelectric Bimorph Bending Actuator with Wires, Thorlabs) was used to perform the whisker pad stimulation protocol (1s stimulation at 3600 deg/s with 20s inter-stimulus interval, for a total of 40-50 trials). To allow simultaneous stimulation of all the contralateral whiskers, they were glued together using a small amount of super-glue (Loctite), before placing the animal in the stereotaxic frame. This stimulation protocol was chosen to mimic natural, rhythmic patterns of whisker activity (Cao et al., 2012; Mégevand et al., 2009).

### Analysis of *in vivo* data

The cortical depth of each recording was evaluated using current-source density (CSD) maps, calculated as the second spatial derivative of the local field potential (LFP) averaged across all the trials. The shortest latency CSD sink evoked upon sensory stimulation was used to define the granular layer. Channels above and below that were assigned to supragranular and infragranular layers respectively. LFP traces were filtered with a 500 Hz low pass filter. Recordings with more than 50% of the power in the noise frequency (∼50Hz) were excluded from further analysis. To analyze spontaneous activity, LFP power spectra were computed using a fast Fourier transform focusing on frequencies up to 20Hz.

Prior to analysis, recordings were spike sorted with Kilosort (Pachitariu et al., 2016) and inspected in Phy (Rossant et al., 2016) to detect multi-unit activity (MUA). Spontaneous MUA activity was evaluated by measuring the average spike rate within each layer during a 20-minute baseline period. To analyze whisker-evoked MUA, spike rate was averaged across trials. MUAs identified within the same layer were then averaged together. The peak response was identified within a 50ms window after the first whisker deflection as a response 3 times bigger than the standard deviation of the baseline activity (evaluated as the mean activity 50ms before stimulation). Whisker-evoked firing rate was evaluated as the difference between firing rate at baseline and maximal firing rate after stimulation of all the layer-averaged MUAs.

### Statistical Analysis

All results in the text are expressed as mean ± standard deviation of the mean; n indicates the number of cells recorded in each independent experiment, unless otherwise stated. In the boxplots, circles represent single data points, horizontal line the median of the data, the box borders indicate the 25th and 75th percentiles and error bars the spread of the data. Microsoft Excel (2010, Microsoft, UK) was used for data organization, while statistical analysis was performed with GraphPad Prism version 6.0 (GraphPad) and with Matlab 2018a (Mathworks). Fisher’s exact test was used to compare synaptic connection incidence. Continuous data were assessed for normality with the D’Agostino-Pearson normality test and for equal variance, in order to apply the appropriate parametric (ANOVAs, t-test) and non-parametric tests (Kruskal–Wallis ANOVAs, Mann–Whitney U test). Statistical significance was evaluated at p≤0.05. Further details of the analysis can be found in the Supplementary Table 2 – Statistics.

## AUTHORS CONTRIBUTION

CV and SJBB designed the research and wrote the manuscript. CV, LJB, FG and SR conducted experiments and analyzed the data. ZM provided mentorship and supervision to CV and access to histological equipment. All authors edited the manuscript.

## ACKNOWLEDGEMENTS

Research in the Butt lab was funded by the BBSRC (BB/P003796/1). CV was funded by the Clarendon Fund and Christ Church College Joint Award in conjunction with a Schorstein Research Fellowship and the Medical Research Council Fellowship. SR was part funded by a small Research Grant from Keble College awarded to SB. FG was funded by a Wellcome Trust DPhil scholarship (215199/Z/19/Z).

We would like to thank Prof. Fumio Matsuzaki (RIKEN Center for Biosystems Dynamics Research, Kobe, Japan) for his generous gift of the conditional *Prox1* mouse line; Prof. Arruda-Carvalho (Department of Psychology, Toronto, Canada) for sharing her experience with early postnatal viral injections; Drs Michael Kohl and Ana Bottura de Barros (Oxford, UK) for their help and training in postnatal injections; and the Micron Advanced Bioimaging Unit (supported by Wellcome Strategic Awards 091911/B/10/Z and 107457/Z/15/Z) for their assistance in this work. We would also like to thank Drs. Adam Packer and Armin Lak for providing feedback on an earlier version of the manuscript.

## SUPPLEMENTARY MATERIAL

**Supplementary Figure 1.**
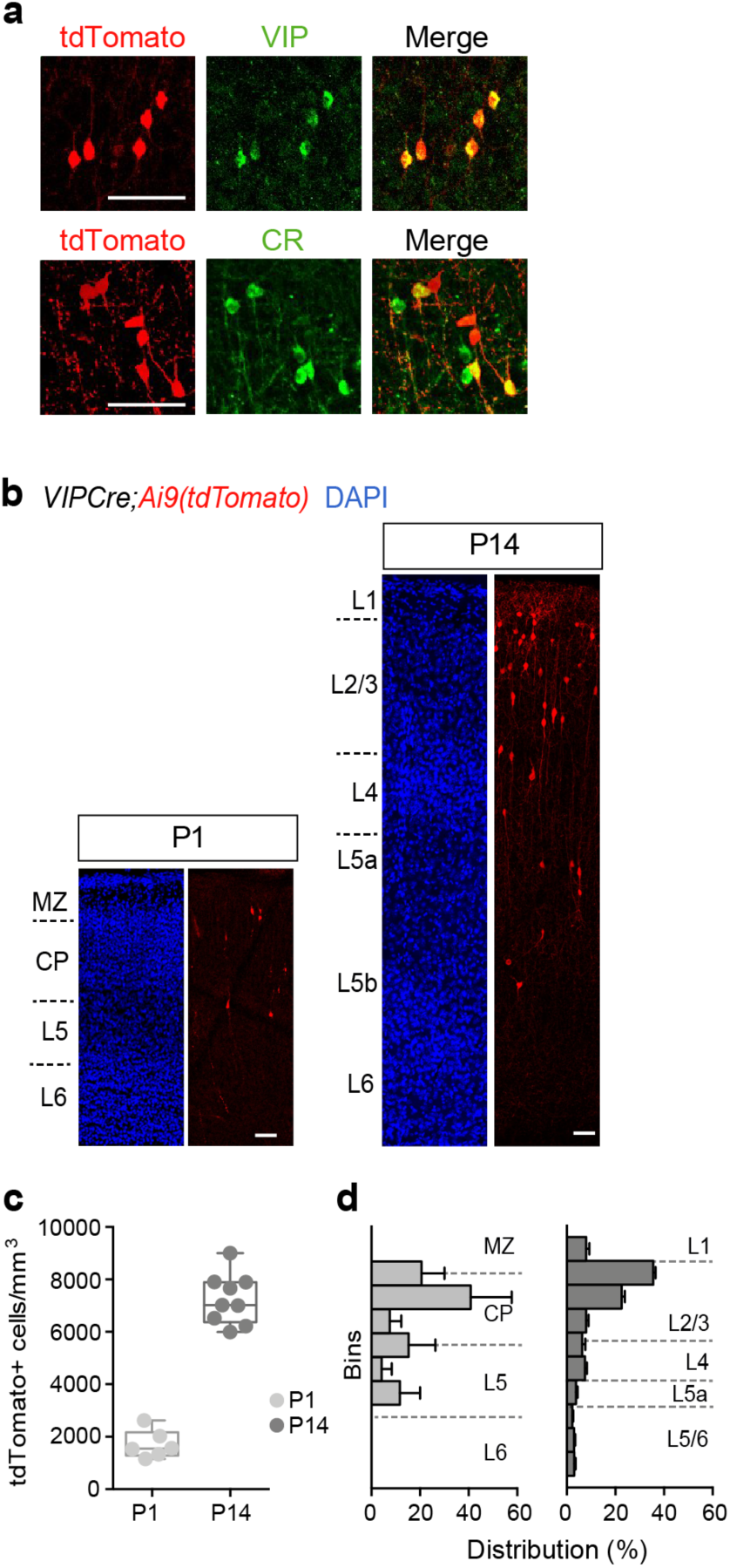
Additional characterization of VIP+ INs in early postnatal S1BF. (a) Images of *VIP-Cre;Ai9* P21 cortex demonstrating co-expression of tdTomato with VIP (top row) and calretinin (bottom); scale bar, 50µm. (b) Images across the depth of cortex in *VIP-Cre;Ai9* mice at postnatal days (P)1 and P14. Scale bar 50µm. MZ, marginal zone; CP, cortical plate. (c) Density of tdTomato+ cells at P1 and P14. (d) Normalized distribution of tdTomato+ cells across the depth of cortex at these two time points. Bars represent the mean ±SEM.

**Supplementary Table 1.**
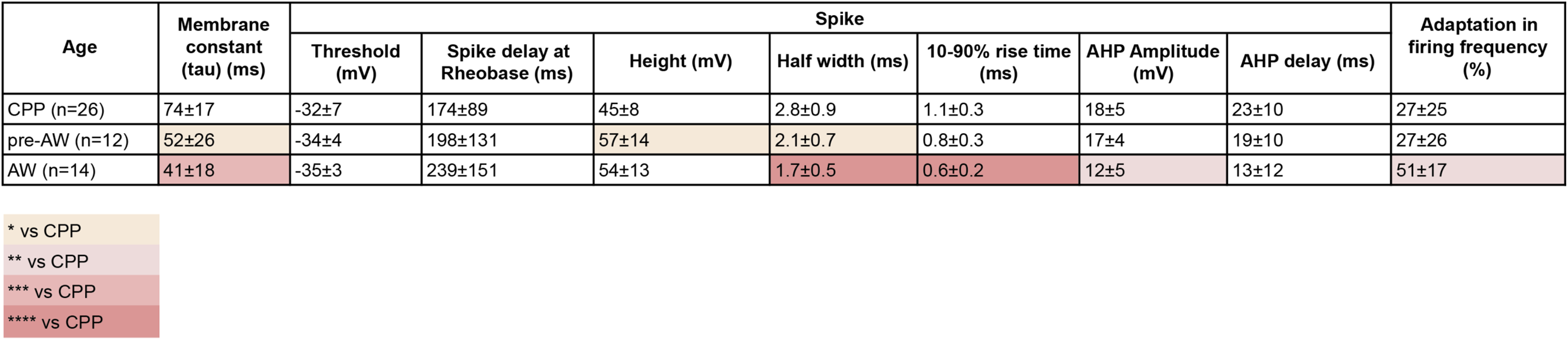
Intrinsic electrophysiological properties of VIP+ INs across early postnatal development. All data is reported as mean ± SD. 10-90% rise time, time from the 10% to the 90% of the action potential height; AHP, after hyperpolarization. Multiple comparisons corrected results following one-way ANOVA or Kruskal Wallis one-way ANOVA are color-coded according to the legend. .*=P<0.05; **=P<0.01; ***<0.001; ****=P<0.0001

**Supplementary Figure 2.**
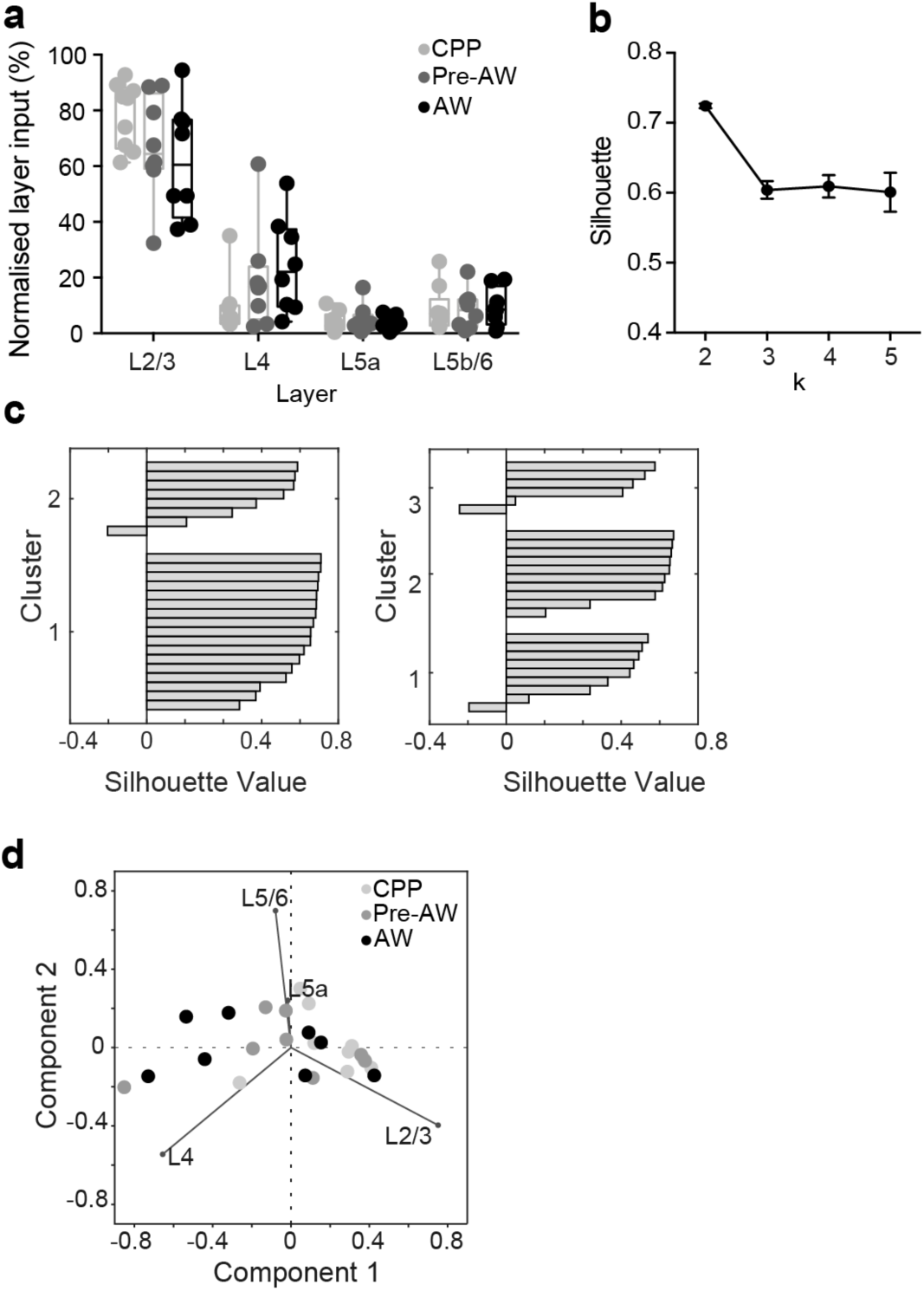
Analysis of LSPS glutamatergic input profiles for VIP+ INs. (a) Percentage of glutamatergic input from each layer onto L2/3 VIP+ INs across postnatal development. (b) Average silhouette values following k-means analysis (1000 repetition per number of clusters) for 2 to 5 clusters (k); k=2 is the optimal number of clusters for classifying glutamatergic afferent input profiles onto VIP+ INs across the time window studied. (c) Silhouette analysis on k-means clustering from k=2 and k=3. (d) Scatter plot of the first two principal components scores following principal component analysis (PCA) of normalized glutamatergic layer input of VIP+ INs. Developmental time points are indicated by different colors.

**Supplementary Figure 5.**
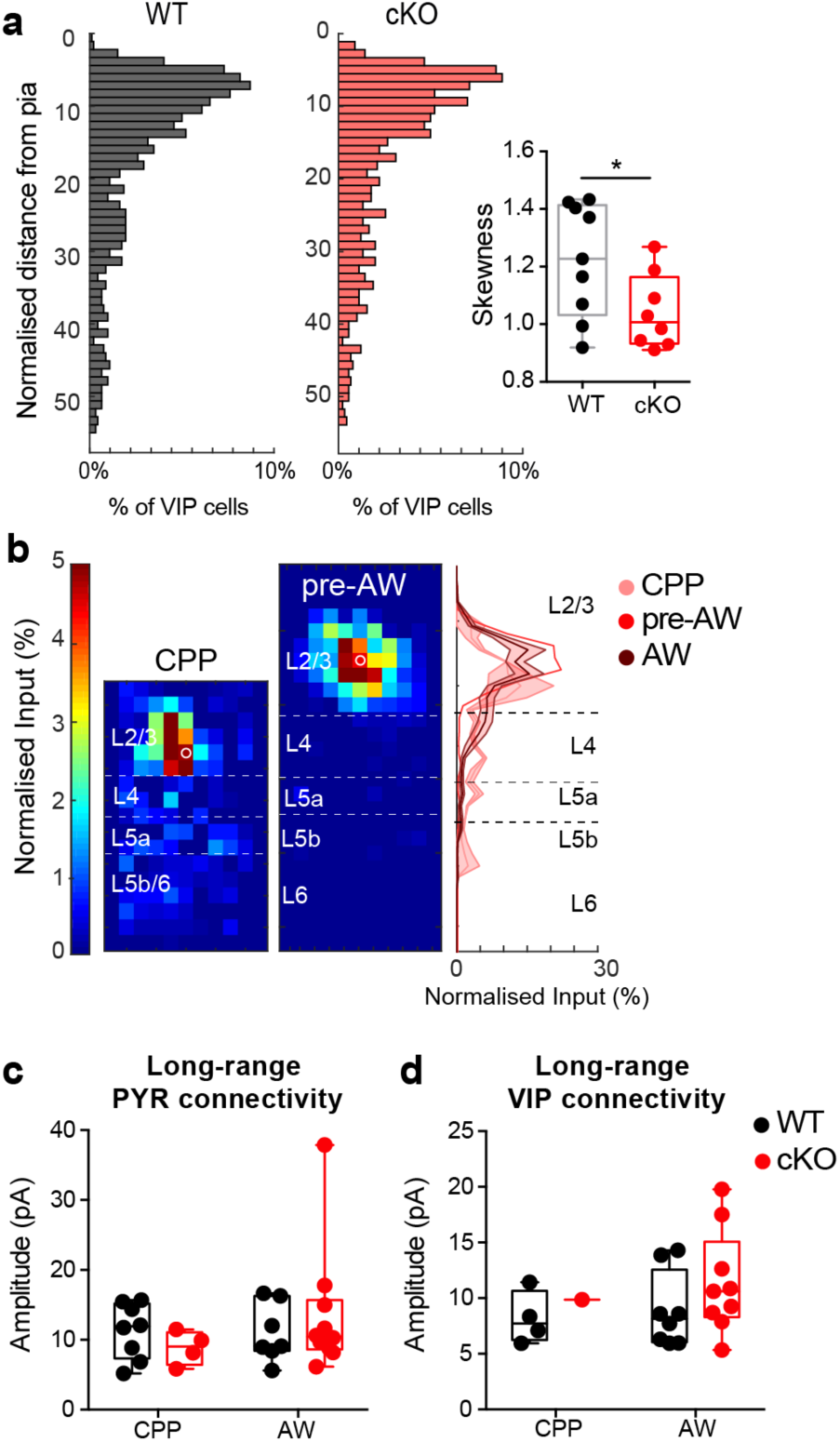
Impact of cell autonomous deletion of *Prox1* on VIP+ interneurons. (a) Distribution of VIP+ INs across S1BF in WT and cKO animals. (Right) Bar plot of skewness (i.e. measure of asymmetry) of the distribution of VIP+ cells in WT vs cKO S1BF. WT: 9 counts from n=3 animals; cKO: 8 counts from n=3 animals. Two tailed, unpaired t test with equal SD * P ≤ 0.05. (b)(left) LSPS maps showing normalised glutamatergic input onto L2/3 cKO VIP+ INs cells recorded prior to AW. (right panel) Layer profile of cKO cells across the three developmental time windows. (c,d) No differences were found in amplitude of long-range inputs from anterior-motor areas onto PYR (c) and VIP+ INs (d) between WT and cKO at any time point.

